# Multiple new species of *Ophiocordyceps* fungus on ants

**DOI:** 10.1101/017723

**Authors:** João P. M. Araújo, David P. Hughes

## Abstract

In tropical forests, one of the most common relationships between parasites and insects is that between the fungus *Ophiocordyceps* (Ophiocordycipitaceae, Hypocreales, Ascomycota) and ants, especially within the tribe Camponotini. These fungi have the ability to penetrate the exoskeleton of the ant and to manipulate the behavior of the host, making it leave the nest and ascend understorey shrubs, to die biting onto the vegetation: hence, the term zombie-ant fungi to describe this behavioral changes on the host. It is posited that this behavioral change aids spore dispersal and thus increases the chances of infection. Despite their undoubted importance for ecosystem functioning, these fungal pathogens are still poorly documented, especially regarding their diversity, ecology and evolutionary relationships. Here, we describe multiple new and host-specific species of the genus *Ophiocordyceps* on *Camponotus* and *Polyrhachis* ants from the central Amazonian region of Brazil, USA, Australia and Japan, which can readily be separated using classic taxonomic criteria, in particular ascospore morphology.

## Introduction

Fungi associated with insects are one of the most spectacular and diverse interactions found on nature. There is an enormous variety to consider: mutualistic symbionts (Suh et al. 2005), fungi serving as an obligate food source, such as those found in fungus-gardening ants (Currie et al. 2003), sexually and behaviorally transmitted parasites – e.g. Laboulbeniales (DeKesel 1996) – and pathogens that have pronounced effects on host populations, termed entomopathogens (Evans 1974). Despite this knowledge, fungal-insect associations remain an understudied area of fungal biodiversity and likely harbor one of the largest reservoirs of undocumented species.

Insects, with over than a million described species (Foottit and Adler 2009), are distributed among 30 orders (sensu Grimaldi & Engel 2005). The fungal pathogens are able to colonize 19 of them, exhibiting a unique diversity of morphologies and strategies to infect those hosts, which also exhibit a myriad of ecologies (Araújo & Hughes in prep.). Among those, one of the most impressive and sophisticated interactions between insects and entomopathogenic fungi is the one between ants and *Ophiocordyceps* fungi (Andersen et al. 2009 life of dead).

Ants are the most abundant land-dwelling arthropods with close to 13,000 species described (Agosti & Johnson 2009). These colonial insects occupy a wide range of habitats, from high canopy to the leaf litter, forming colonies comprising from a couple of dozens to millions of individuals, especially on tropical forests. However, inhabit such a prolific environment is a tradeoff.

*Ophiocordyceps* fungi, a predominantly entomopathogenic genus – comprised by about 160 species (Sung et al. 2007) – is also very well adapted to live in the warm and humid tropical forests. The genus emerged about 100 mya (Sung et al. 2008) and since then is infecting ten orders of insects. Among those, a specific group is notable to be able to trigger a sophisticated synchronized chain of events that will lead the host to die in an exact position for the fungal optimal development. Those are the species belonging to a complex of species, named *Ophiocordyceps unilateralis sensu lato.* They are able to lead the infected ants to leave the colony and climb up to the underside or edge of leaves, locking its jaws into the plant tissue to ensure the proper attachment for the subsequent weeks of fungal development. About two weeks later, the fungus has already developed the fruiting body (ascoma) to release their infectious units (spores) actively on the forest floor.

Since *Ophiocordyceps unilateralis* was originally published as *Torrubia unilateralis* (Tulasne & Tulasne 1865), few species were described belonging to the *O. unilateralis s.l.* group so far (Evans & Samson 1984; Evans et al. 2011; Kepler et al. 2011; Luangsa-ard et al. 2011; Kobmo et al. 2012), despite their diversity being estimated for about 580 species worldwide (Araújo & Hughes, unpublished data). In addition, Kobayasi (1939; 1982) described a different species as being a different variety (*O. unilateralis* var. clavata) from Japan, which we propose to be erected as a new species rather than variety herein.

## Material and methods

### Sampling

Surveys were undertaken in the central Amazonian region of Brazil at Reserva Adolpho Ducke. The reserve is comprised by ca. 10,000 ha (02°55’S, 59°59’W), adjacent to Manaus (Amazonas state) and composed of terra-firme forest, with plateaus, lowlands and campinarana vegetation, characterized to occur in patches of sandy soil across the Rio Negro basin.

Sampling protocol consisted of a careful inspection of soil, leaf litter, shrub leaves and tree trunks, up to *ca.* 2 m high. Infected ants – and the substrata they were attached to – were collected in plastic containers for transport to the laboratory and, wherever possible, examined the same day. During longer surveys, the samples were air-dried overnight to prevent mold growth. All specimens were photographed individually, using a Canon 60D camera fitted with a MP-E 65mm (x5) macro lens, equipped with a MT-24EX macro lite flash.

### Morphological studies

Samples were screened using a stereoscopic microscope, and only mature fungal specimens were selected for further micro-morphological studies. In order to obtain ascospores, infected ants were attached to the lid of a plastic Petri dish using petroleum jelly, and suspended above a plate containing either distilled-water agar (DWA) or potato dextrose-agar (PDA). Plates containing the ants attached were maintained outside the lab at natural temperature and examined daily for the presence of ascospores, which, after ejection from the ascoma, formed sub-hyaline halos on the agar surface. Freshly-deposited ascospores were removed with a sterile hypodermic needle, with the aid of a stereoscopic microscope, and mounted on a slide in lacto-fuchsin (0.1g of acid fuchsin in 100 ml of lactic acid) for light microscopy (Olympus BX61). A minimum of 50 naturally-released ascospores were measured for morphological comparison (Table 1). The remaining ascospores were left *in situ* on the agar surface and examined over a number of days in order to monitor germination events. For micro-morphology of the ascomata, either free-hand or cryo-sectioning (Leica CM1950 Cryostat) was used.

## Results

### Taxonomic treatment

***Ophiocordyceps briophyticola*** Araújo & D. P. Hughes sp. nov.

External mycelium covering most of the host, produced from all orifices and sutures, brown at maturity. Stroma single, rarely branched, produced from dorsal pronotum, averaging 15–20 mm, up to 30 mm, cylindrical, velvety and dark brown, tapering towards the apex; Fertile region (ascoma) of lateral cushion, 1-2, hemispherical to globose, dark-brown to black, variable in size, averaging 1-1.5 x 0.8-1 mm. Perithecia immersed to partially erumpent, flask-shaped, 220–250 x 100–165 µm, pronounced ostiole. Asci 8-spored, hyaline, cylindrical, (110-) 130-145 x 8 – 10µm; prominent cap, 7–8 x 3 µm. Ascospores hyaline, thin walled, vermiform 90 – 120 x 4 µm, 5–8-septate, straight to sinuous, round to slightly tapered apex.

**Asexual-morph.** Hirsutella A-type not observed. Hirsutella C-type, produced from brown cushions (sporodochia) on leg and antennal joints; phialides subulate at base, 40–60 x 3–5 µm long, tapering to a long, hyaline neck. Conidia not observed.

**Germination process.** All the ascospores remained unchanged after five days on water-agar plate. Further studies will address more attempts to incubate the ascospores for more days, aiming hyphal or capilliconidiophore germination.

**Figure 1.**
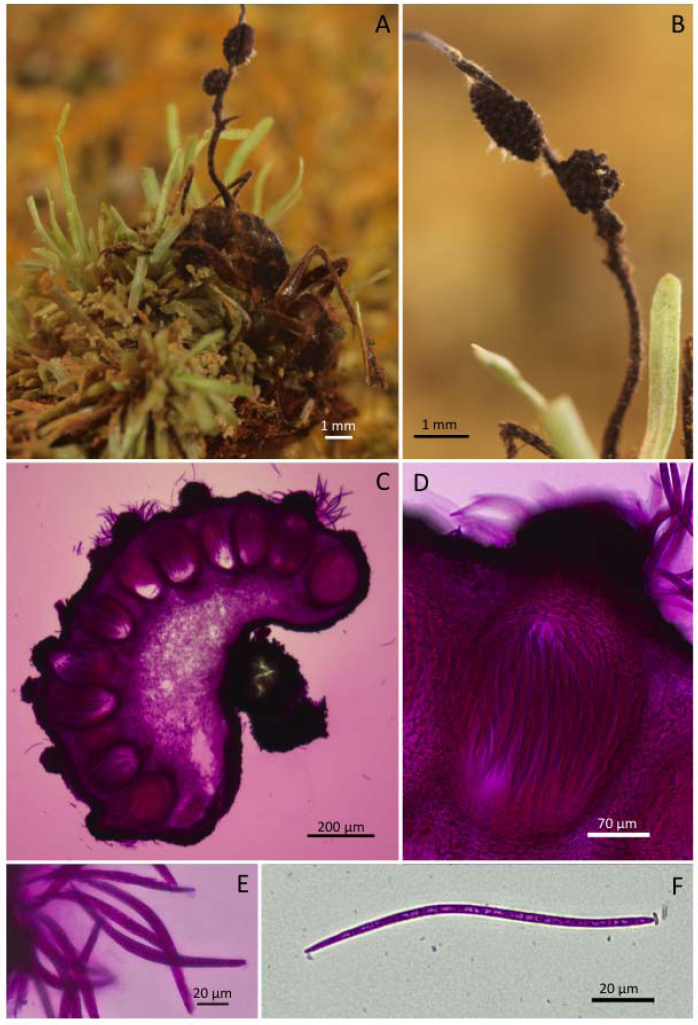
Ophiocordyceps bryophyticola. **A)** *Camponotus* sp. dying attached to bryophytes on the base of trees; **B)** Close-up of the fertile part (ascoma); **C)** Section through ascoma showing the perithecial arrangement; **D)** Close-up of perithecium; **E)** Asci; **F)** Ascospores

**Figure 2.**
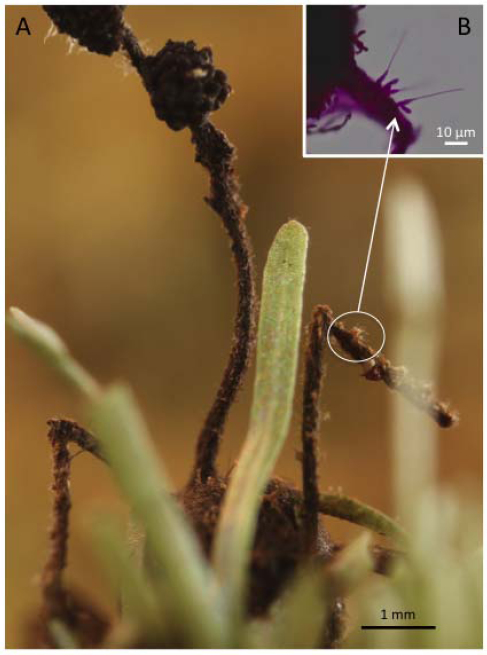
Ophiocordyceps bryophyticola. **A)** Ant biting into the moss carpet with the antenna raised, showing the detail of **B)** phialides (Hirsutella B-type).

***Ophiocordyceps chantifex*** Araújo & D. P. Hughes sp. nov.

Mycelium growing from all inter-segmental membranes, often covering the host body; initially white turning brown. Stroma single, produced from dorsal pronotum, averaging 10 mm, up to 15 mm in length, cylindrical, velvety and ginger brown, becoming cream-pinkish at the apical part; fertile region of lateral cushion, 1–2, hemispherical, chocolate brown, darkening with age, slightly variable in size, averaging 1.5 x 1 mm. Perithecia immersed to partially erumpent, globose-hemispherical shaped, 200–235 x 135–175, with short neck. Asci 8-spored, hyaline, cylindrical to clavate, 100–125 x 6 µm; prominent cap, 7–6 x 3–4 µm. Ascospore hyaline, thin-walled, vermiform 75–85 x 5 µm, 9–13-septated, sinuous to curved, never straight at maturity, rounded to acute apex.

**Asexual-morph.** Hirsutella-A type associated with apical region of stromata; phialides lageniform, 5–6 x 3 µm, tapering to a robust neck, 4–8 µm in length; conidia fusiform to limoniform, averaging 7 x 2.6 µm.

**Germination process.** The released ascospores germinated within 24h to produce a single, long and extremely narrow hair-like capilliconidiophore; variable in length (65-) 75–90 (−95) µm; bearing a single terminal capilliconidium, hyaline, smooth-walled, uni or biguttulate, fusoid, narrowing apically.

**Figure 3.**
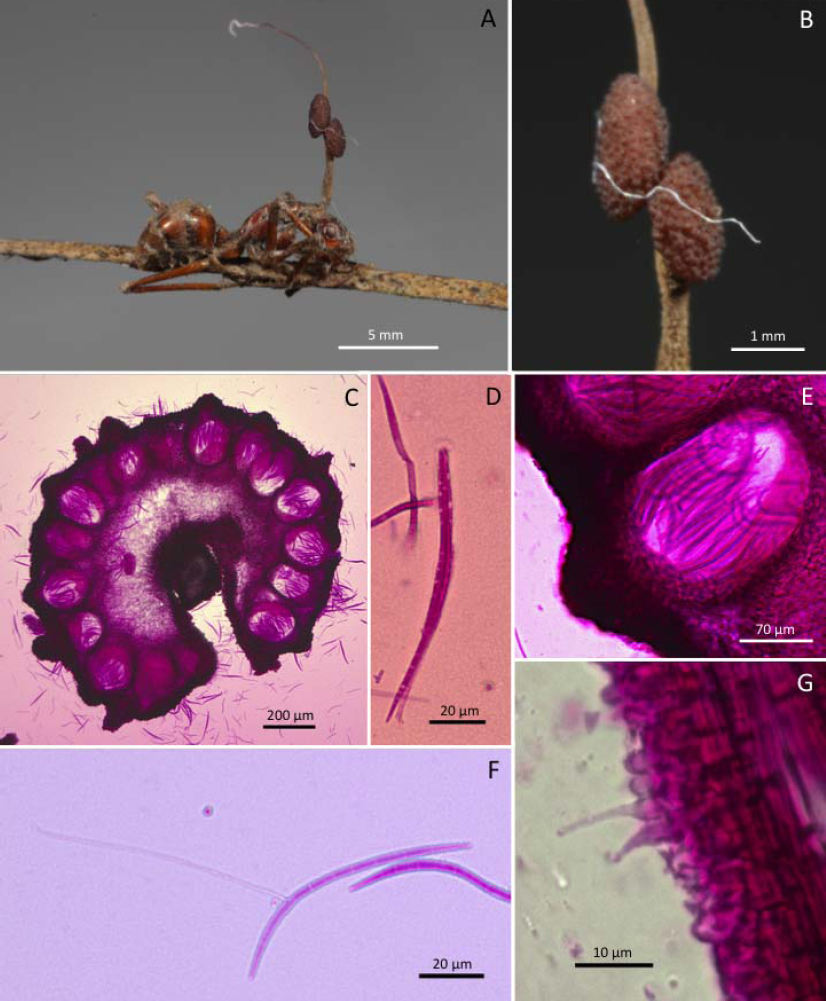
Ophiocordyceps chantifex. **A)** Camponotus chantifex biting onto a palm leaf; **B)** Close-up of the ascoma; **C)** Cross section of the ascoma showing the perithecial arrangement; **D)** Ascus with prominent cap; **E)** Close-up of the perithecium; **F)** Multiseptated ascospore with long capilliconidiophore; **G)** Hirsutella A-type phialide on the stroma.

***Ophiocordyceps elegans*** Araújo & D. P. Hughes sp. nov.

Mycelium produced sparsely from joints, not covering the host body, dense when touching the substrate, dark brown. Stroma single, arising from the dorsal pronotum, never branching, averaging 1.8−2 cm in length, 0.2 mm thick, dark brown at the base turning into lighter brown into the apex; fertile part of a single lateral cushion, disc-shaped, chestnut-brown, averaging 1x1 mm. Perithecia immersed to partially erumpent, flask-shaped, (205-) 225–230 (−265) x 135 (−180) µm with short neck. Asci 8-spored, hyaline, cylindrical, 150–160 x 8–9 µm; apical cap prominent, 6 x 3 µm. Ascospores hyaline, thin-walled, multiguttulate, cylindrical, 120–140 x 3µm, 7-septate, straight or curved tapering to the apex.

**Anamorph.** Hirsutella A-type associated with apical region of stroma; phialides lageniform, 5–8 x 3–4 µm, tapering to a long neck, 8–12 µm; conidia hyaline, limoniform, 5x2 µm.

**Germination process.** Ascospores released on agar germinated after 72h to produce a single, straight capillicondiophore; 25–30 µm, bearing a terminal capilliconidium, hyaline, smooth-walled, guttulate, 5–9 x2 µm, narrowing apically.

**Figure 4.**
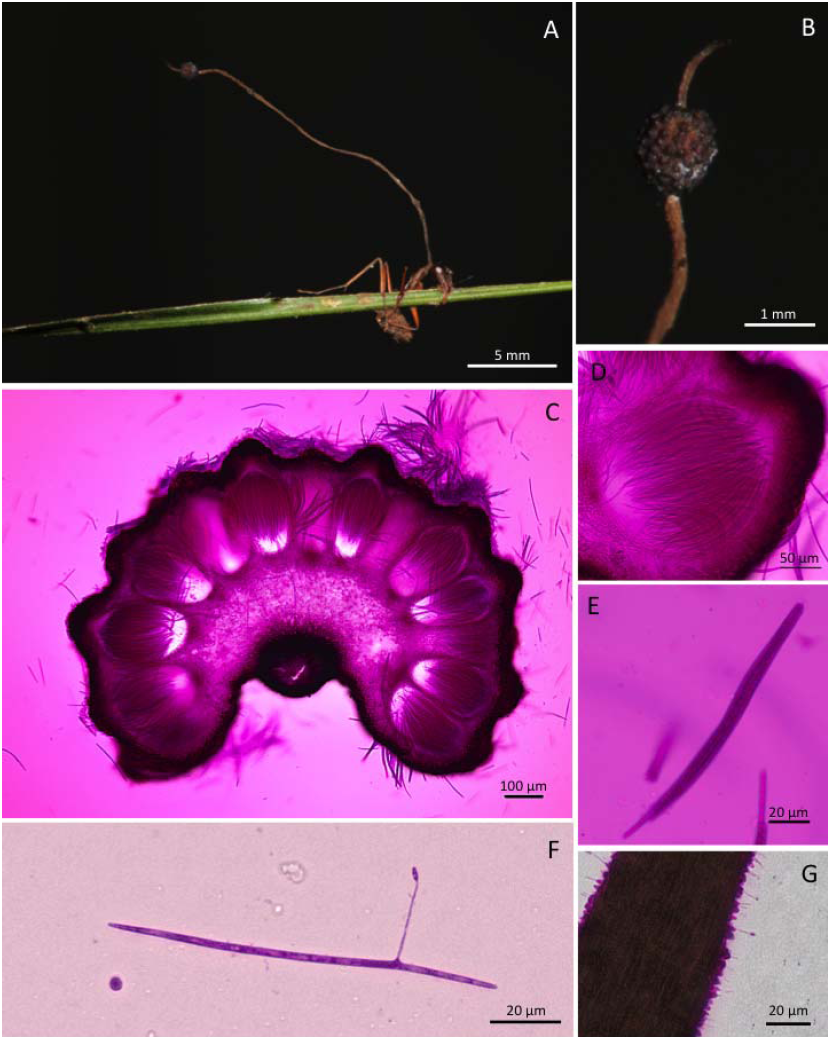
Ophiocordyceps elegans. **A)** *Camponotus* sp. biting into vegetation with the long stroma arising from its dorsal pronotum; **B)** Close-up of the ascoma; **C)** Section through ascoma showinh the perithecial arrangement; **D)** Close-up of perithecium; **E)** Ascus; **F)** Long ascospores with the straight capilliconidiophore bearing an apical capilliconidium; **G)** Hirsutella A-type layer on the apical part of the stroma.

***Ophiocordyceps sporodochialis-nigrans*** Araújo & D. P. Hughes sp. nov.

External mycelium produced from all orifices and sutures; initially white, becoming ginger brown, covering the host body, notably the abdominal part. Stroma single, produced from dorsal pronotum, 10–15 x 0.2 mm, cylindrical, black, covered with ginger velvety hyphae fading away towards the apex; fertile region of lateral cushions, 1–2, disc-shaped to hemispherical, light brown, darkening with age, averaging 1.5 x 1 mm. Perithecia immersed to partially erumpent, flask-shaped, 215–240 x 120–150 (−180) µm, with short, exposed neck or ostiole. Asci 8-spored, hyaline, thin-walled, vermiform to clavate, 120–145 x 8 (−10) µm; cap prominent, 8 x 4 µm; Ascospores hyaline, thin-walled, vermiform, 90–105 (−115) x 3−4 µm, 5-septate, gently curved, rarely straight; tapering to a round apex.

**Anamorph.** Hirsutella A-type associated with the apical part of stroma. Hirsutella C-type, produced from light brown cushions on leg and antennal joints: phialides subulate, robust, 85–120 x 4–6 (−8) µm.

**Germination process.** Ascospores germinating after 24–72h to produce 1 (−2), uniformly straight, extremely narrow hair-like capilliconidiophores, 50–60 µm; bearing a single terminal capilliconidium, hyaline, smooth-walled, biguttulate, clavate, 9 x 2 µm, narrowing apically.

**Figure 5.**
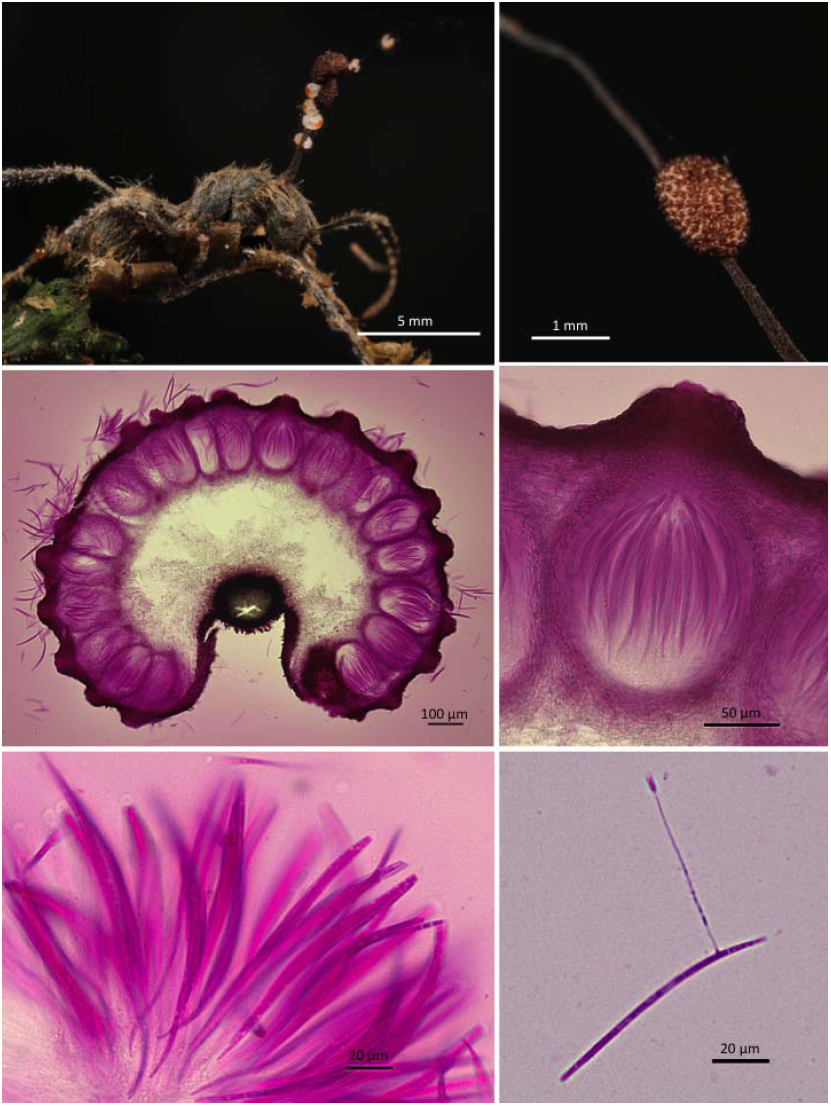
Ophiocordyceps sporodochialis-nigrans. **A)** *Camponotus* sp. infected biting into a leaf; **B)** Close-up of the ascoma; **C)** Section through ascoma showing the perithecia arrangement; **D)** Close-up of perithecium; **E)** Asci; **F)** Ascospore with capilliconidium.

**Figure 6.**
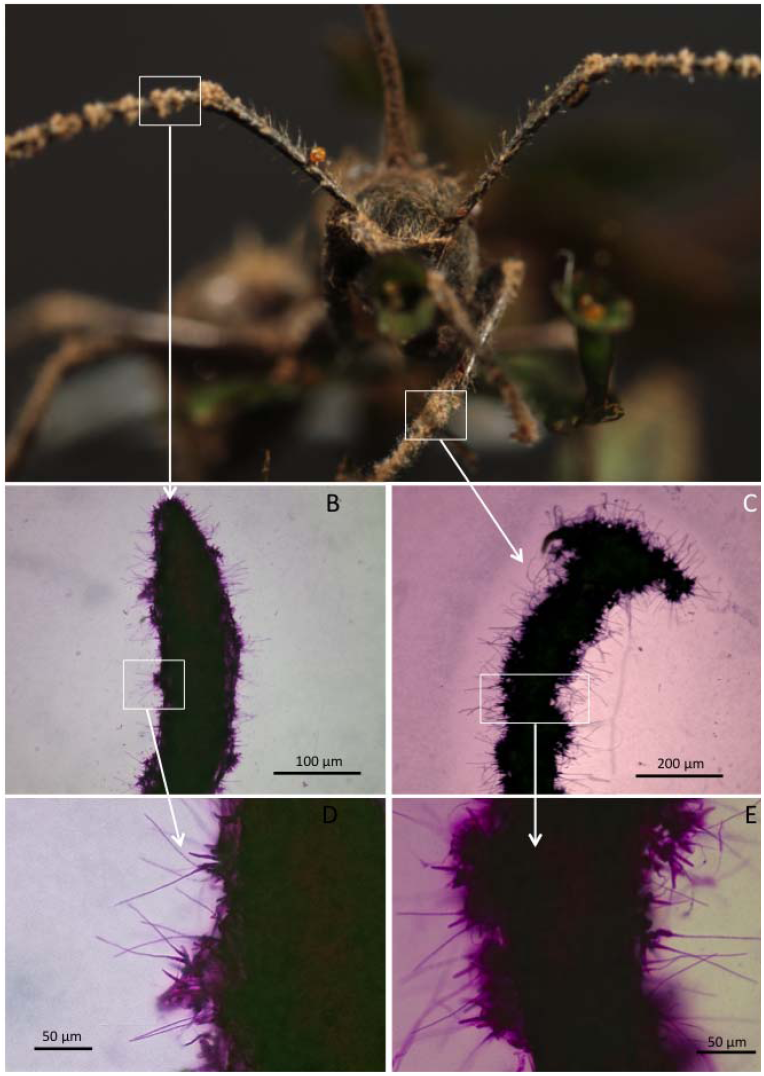
Ophiocordyceps sporodochialis-nigrans. Plate showing the Hirsutella C-type growing on legs and antenna. A) Frontal view of the biting ant; B) Antenna covered with phialidesl C) eg covered with phialides; D-E) Close-up of the subulate phialides.

***Ophiocordyceps sporodochialis-brunneis*** Araújo & D. P. Hughes sp. nov.

External mycelium produced from all the orifices and sutures; initially white, becoming ginger brown, covering the host body with sparse hyphae. Stroma single, produced from dorsal pronotum, averaging 15 x 0.2 mm, cylindrical, black, covered with ginger velvety up to the ascoma, fading away towards the apex; fertile region of a single lateral cushion, disc-shaped, chestnut-brown, darkening with age, averaging 1.3 x 1 mm. Perithecia immersed to partially erumpent, flask-shaped, (170-) 200−215 x 100–130 µm, with short, exposed neck or ostiole. Asci 8-spored, hyaline, thin-walled, vermiform to clavate, 110–140 x 6–8 µm; cap prominent, 4x6 µm; Ascospores hyaline, thin-walled, vermiform, 75–90 x 3 µm, 5-septate, straight to gently curved, tapering to a round apex.

**Anamorph.** Hirsutella A-type associated with the apical part of stroma. Hirsutella C-type, produced from light brown cushions on leg and antennal joints: phialides subulate, robust, 70–95 x 5 µm.

**Germination process.** Ascospores germinating after 24–72h to produce 1–3, extremely narrow hair-like capilliconidiophores, 50–60 µm; bearing a single terminal capilliconidium, hyaline, smooth-walled, bi-pluri guttulate, clavate, 9–10 x 3 µm, narrowing apically.

**Figure 7.**
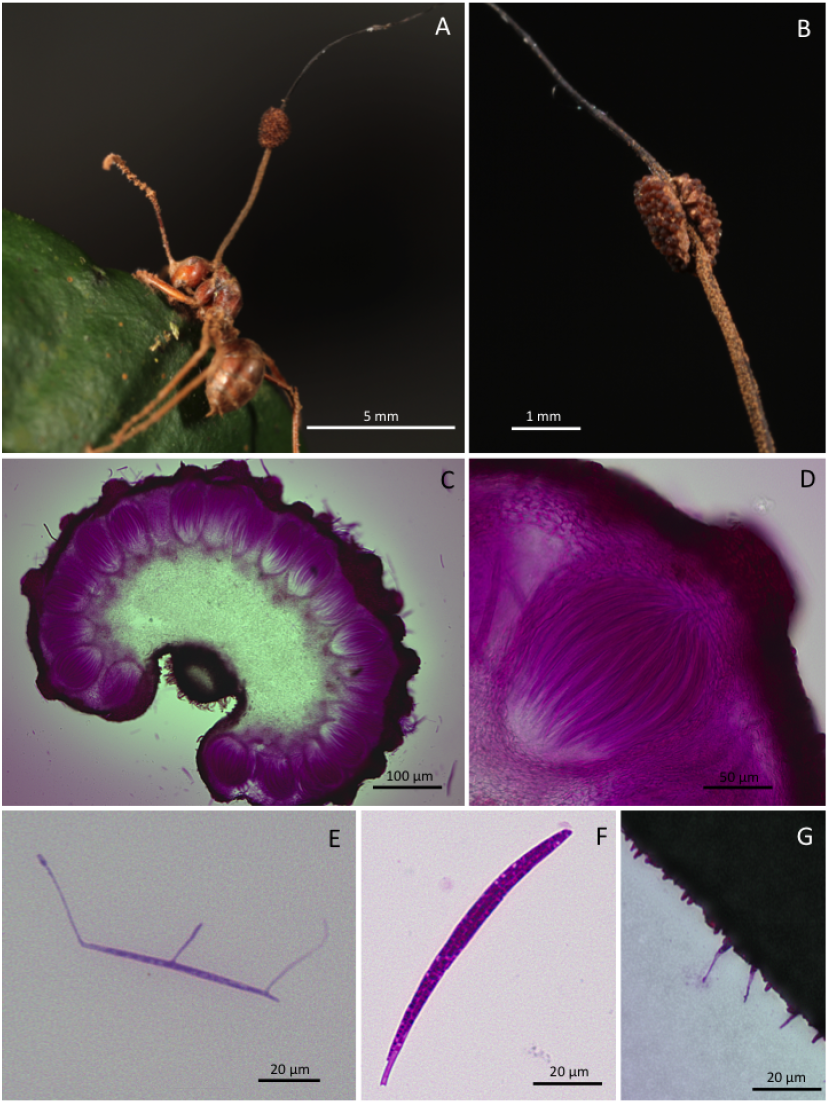
Ophiocordyceps sporodochialis-brunneis. Camponotus sp. biting into a sapling’s leaf edge. **B)** Close-up of the ascoma; **C)** Cross section of the ascoma, showing the perithecial arrangement; **D)** Close-up of ferithecium; **E)** Ascospores showing three capilliconidiophores; **F)** Ascus; **G)** Hirsutella A-type phialides.

**Figure 8.**
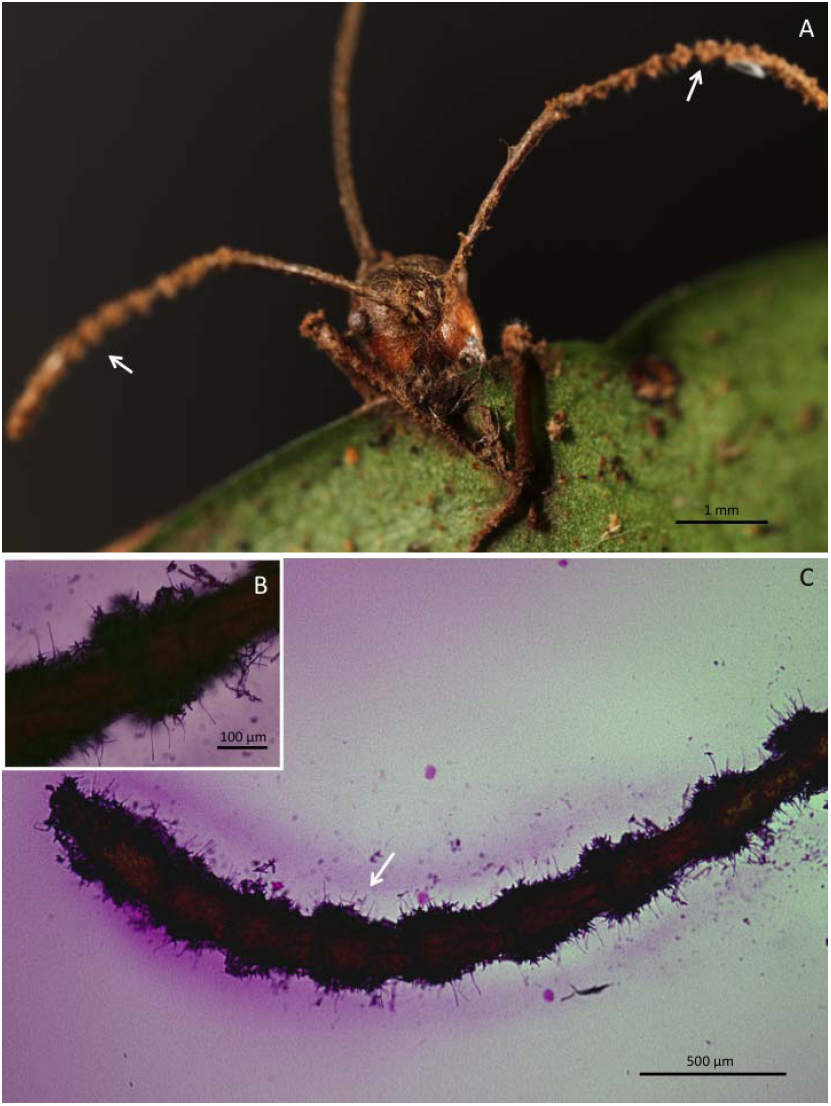
Ophiocordyceps sporodochialis-brunneis. **A)** Frontal view of the ant biting the edge of the leaf. B-C) Antenna covered with Hirsutella C-type phialides.

***Ophiocordyceps bicolor*** Araújo & D. P. Hughes sp. nov.

External mycelium produced from all the orifices and sutures; initially white, becoming ginger brown, covering the host body with sparse hyphae. Stroma single, produced from dorsal pronotum, averaging 3.5 x 0.25, up to 6 mm in length, cylindrical to laterally compressed, ginger to dark-brown; fertile part terminal of lateral cushions, 1–3, disc-shaped to hemispherical, chestnut-brown, darkening with age, 1.2 – 2.2 x 0.8–1.4 mm. Perithecia immersed to partially erumpent, flask-shaped, 200–230 (−250) x 135–165 µm, with short, exposed neck or ostiole. Asci 8-spored, hyaline, cylindrical to clavate, 110–130 x 8–9 µm; cap prominent, 6–3 µm; Ascsopores hyaline, sinuous to curved, rarely straight, 75–90 x 3 µm, 5-septate; apex round to acute.

**Anamorph.** Hirsutella A-type only: produced laterally on upper stroma; phialides rare, cylindrical to lageniform, 7–10 x 3–4 µm, tapering to a long neck, 10–15 µm; conidia limoniform, averaging 7–9 x 3 µm.

**Germination process.** Ascospores germinated in 24−48h to produce a single, narrow capilliconidiophore, 35−40 µm long; bearing a single capilliconidium, hyaline, smooth-walled, uni-biguttulate, clavate, 9–3 µm, narrowing apically.

**Figure 9.**
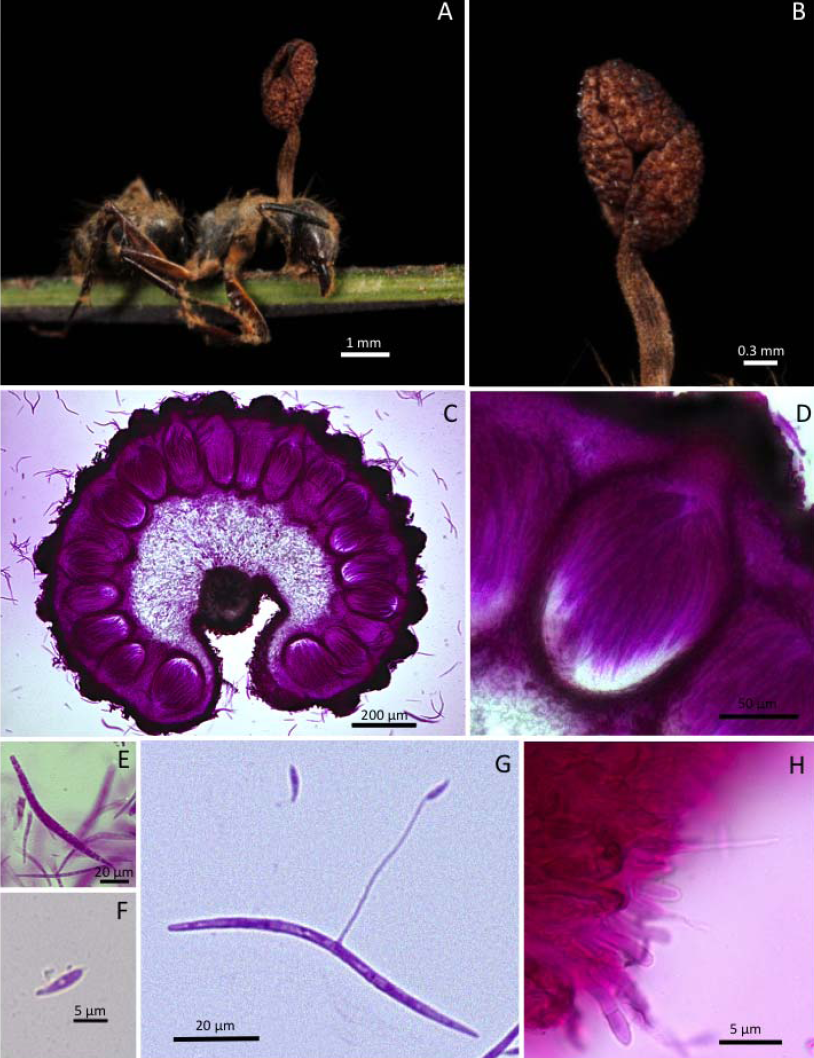
Ophiocordyceps bicolor. **A)** *Camponotus sp* biting a palm leaf; **B)** Close-up of the ascoma; **C)** Cross section show the perithecial arrangement; D) Close-up of the perithecium; E) Ascus; F) Capilliconidium; G) Sinuous ascospore with the capilliconidiophore bearing a capillicondium; H) Hirustella A-type phialide on the stroma.

***Ophiocordyceps triangularis*** Araújo & D. P. Hughes sp. nov.

External mycelium produced from all the orifices and sutures; initially white, becoming ginger brown, covering the host body with sparse hyphae. Stroma single, produced from dorsal pronotum, 5–7 x 0.15 mm, cylindrical ginger to dark-brown, swollen terminal part, clavate; fertile part constantly produced at the middle part of stroma, laterally attached, round-shaped, chestnut brown, darkening with age, averaging 2–2.5 x 0.25–0.45 mm; Perithecia immersed to partially erumpent, flask-shaped, averaging 225–250 x 135–165 µm, with short, exposed neck or ostiole. Asci 8-spored, cylindrical to clavate, 115–135 x 7–10 µm, cap prominent, 6–7 x 4 µm; Ascospores hyaline, cylindrical, robust, straight to gently curved, 75–85 x 4–5 µm, 5-septate, tapering to a round to slightly acute apex.

**Anamorph.** Hirsutella A-type only; produced on the clavate part of upper stromal phialides cylindrical to lageniform, 8–9 x 4 µm, tapering to long neck 9−10 µm long; conidia limoniform, averaging 5 x 2µm.

**Germination process.** Ascospores germinated 24–48h to produce a straight, robust capillicondiophore, often verrucose, 45–50 µm long; bearing a single capilliconidium, hyaline, smooth-walled, guttulate, 10–11 x 4 µm, truncate at base, narrowing apically.

**Figure 10.**
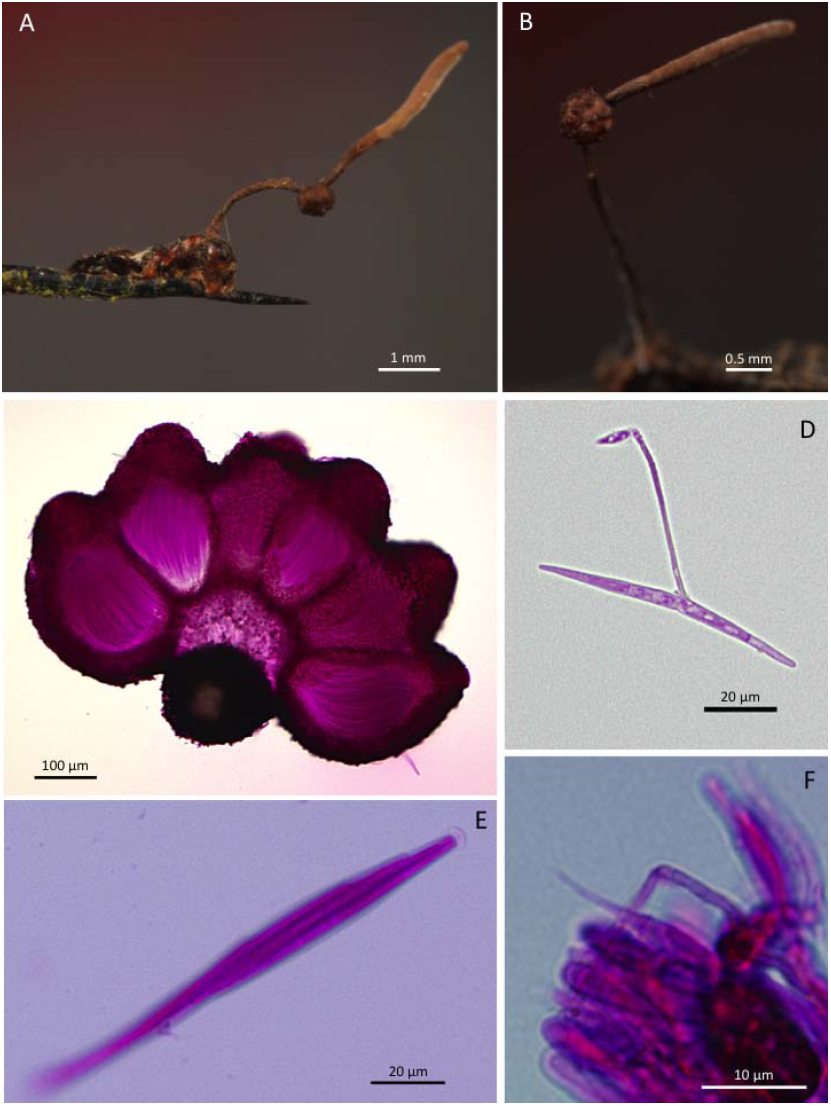
Ophiocordyceps triangularis. **A)** Tiny *Camponotus* sp biting onto a palm spine. **B)** Cloase-up of the ascoma; **C)** Cross section of the ascoma showing the perithecial arrangement; **D)** Ascospore with capilliconidiophore with verrucose apical portion; E) Ascus; F) Hirsutella A-type phialide on the stroma apex.

***Ophiocordyceps terminalis*** Araújo & D. P. Hughes

External mycelium produced from all the orifices and sutures; ginger brown, covering the host body, especially the abdomen. Host body shrunk. Stroma single, produced from dorsal pronotum, 10 x 0.2 mm, cylindrical to flattened dark brown; fertile part disc-shaped, produced terminally on stroma, attached laterally, chestnut to dark brown, 3 x 2.5 mm; Perithecia 235–280 x 130–155 (−180), with short, exposed neck or ostiole. Asci 8-spored, cylindrical to clavate, 110–135 x 6–8, cap prominent, 6x3 µm; Ascospores hyaline, cylindrical, gently curved, 85–95 x 2 µm, 5-septated, tapering to a round, slightly acute apex.

**Anamorph.** Hirsutella A-type only; produced on most of the stroma, phialides cylindrical to lageniform, 4–5 x 3 µm, tapering to a long neck 10–20 µm long, conidia hemispherical, averaging 5x2 µm.

**Germination process.** Ascospores germinated 48–72h to produce one 1–2 straight, short capilliconidiophores, 8–10 µm, bearing a terminal capilliconidium 6–7 x 1–2 µm.

**Fig. 11.**
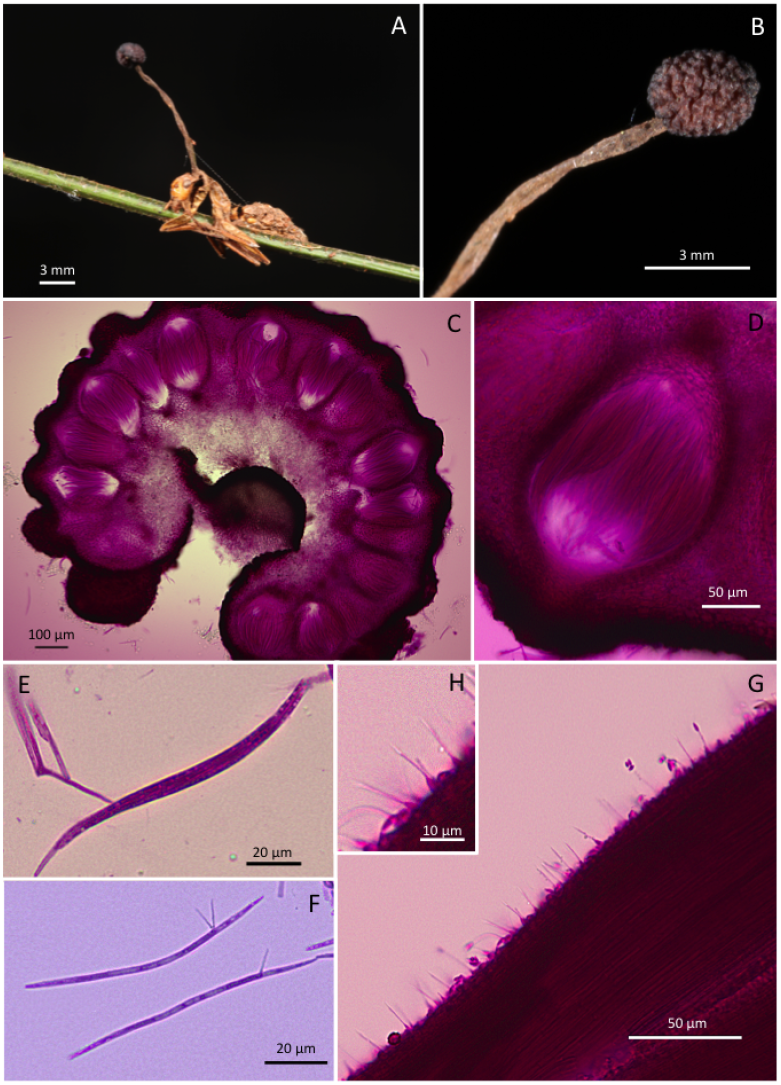
Ophiocordyceps terminalis. **A)** Camponotus sp. biting a palm-leaf; **B)**Close-up of the stroma containing the trminal ascoma; **C)**Perithecial arrangement in a section of the ascoma; **D)** Perithecium with asci inside; **E)** Ascus; **F)**Ascospores with 1–2 capilliconidiophores; **G-H)** Hirsutella type-A on stroma.

***Ophiocordyceps flemmingii*** Araújo & D. P. Hughes sp. nov.

External mycelium produced mostly on the ventral part of the host and head. Sparse mycelium produced on joints. Stroma single, rarely branched, produced from dorsal pronotum, 11–17 x 0.3−.045 mm, cylindrical, ginger to light brown, basal part velvety, apical part cream to purple; fertile part laterally attached, disc-shaped, dark-brown to black, averaging 1.5–2 x 1.3 mm; Perithecia immersed to partially erumpent, flask-shaped, 250−275 x (100-) 120–160 µm, with short, exposed neck or ostiole. Asci 8-spored, cylindrical to clavate, (100-) 120–150 x 10–11 µm, cap prominent; Ascospore hyaline, cylindrical, straight, 80–90 x 5µm, X-septate, tapering towards the apexes;

**Anamorph.** Hirsutella A-type present on the stroma, ; Hirsutella C-type occurring exclusively on early stages of development, produced from leg joints and dorsal pronotum.

**Germination process.** Ascospores germinating from the first 24h up to the 5^th^ day; Germination occurred in two different manners:capilliconidiophores or germination into vegetative hyphae, separately or both on the same ascospores. Capilliconidiophores 1–3, bearing a terminal capilliconidium, 10–13 x 2–3 µm.

**Fig. 12.**
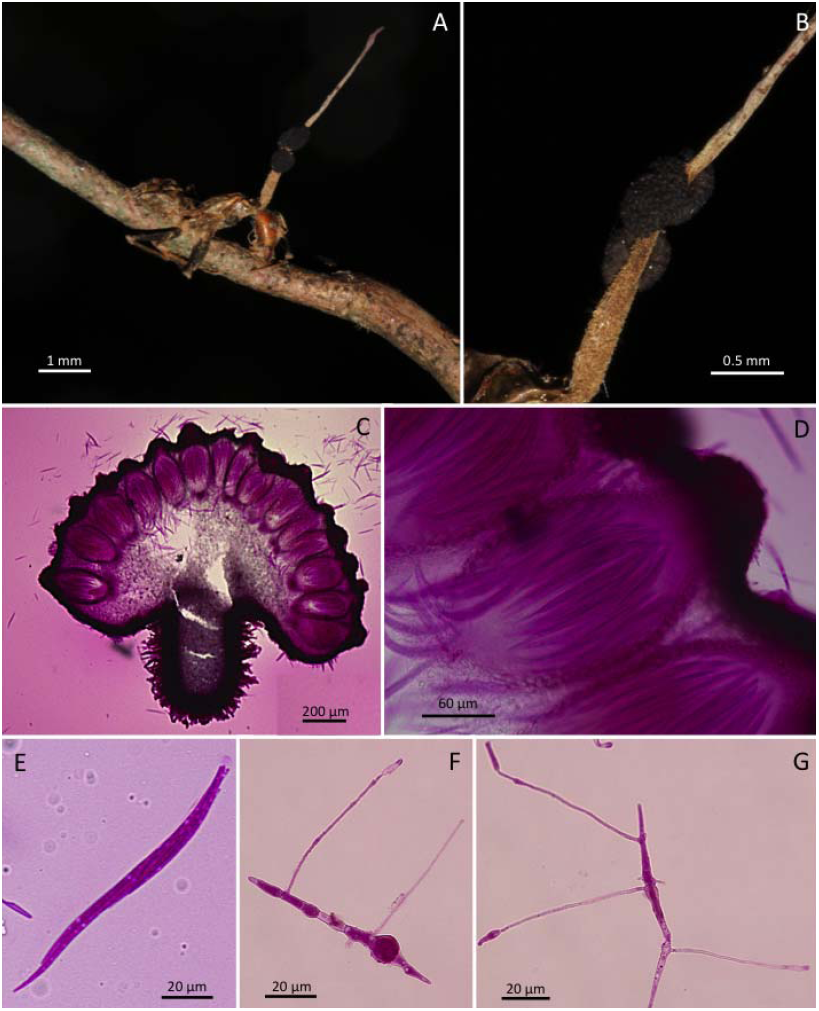
Ophiocordyceps flemmingii (teleomorphic structures). **A)** *Camponotus castaneus* biting a twig; **B)**Close-up of the stroma showing two ascomatal plates attached on it; **C)**Ascoma section and Perithecia arranged on its surface; **D)**Perithecium; **E)**Ascus; **F)**Ascospores after 2–5 days on agar, exhibiting a swollen section and two capilliconidiophores; **G)** Ascospores after 2–5 days on agar showing multiple germination of capilliconidiophores and small hyphal growth on the medium part.

**Fig. 13.**
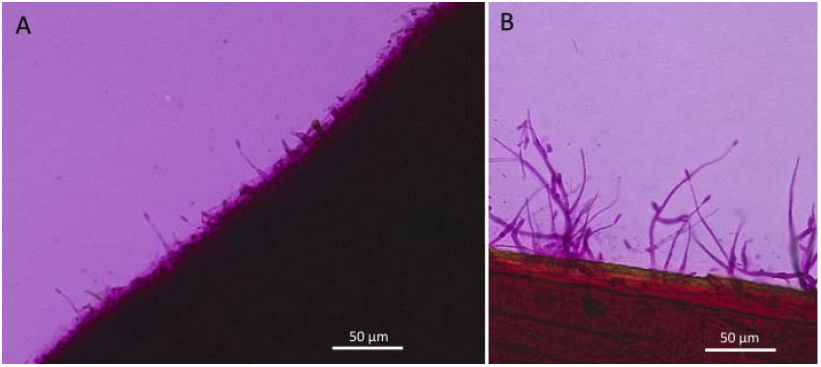
Ophiocordyceps flemmingii (anamorphic structures). **A)** Hirsutella type-A on the stroma; **B)** Hirsutella type-C growing on legs and the very early stage of the stroma, both exclusively on recently dead ants.

***Ophiocordyceps australia*** Araújo & D. P. Hughes sp. nov.

External mycelium produced mostly on the ventral part of the host, scares mycelium present on joints. Stromata ginger to light-brown, commonly clavate, produced always from dorsal pronotum, frequently on leg joints, 1.5–2.25 x 0.45–0.75 mm, branching into nodules formed along the stromata, 120–150 x 35–50 µm, phialides very abundant along the whole stromata; Fertile part single, attached laterally, hemispheric to irregular shape, orange, averaging 0.75 x 0.5–0.65 mm. Perithecia immersed, flask-shaped, 260–320 x (−130) 150–200 µm, with prominent neck. Asci 8-spored, hyaline, vermiform, cylindrical, 150–180 x 7 µm, cap prominent; Ascospore hyaline, straight to gently curved, vermiform, 75–105 x 5–6 µm, 4−6-septate; tapered apex.

**Anamorph.** Hirsutella-X, produced profusely along the whole stromata; phialides abundant, cylindrical to clavate, 15–35 x 3 µm, producing up to 10 needle-like, verrucose conidiophores, averaging 10 µm, bearing a terminal conidium, 5–7 x 3 µm.

**Germination process.** No germination could be observed since the material examined was not alive.

**Fig. 14.**
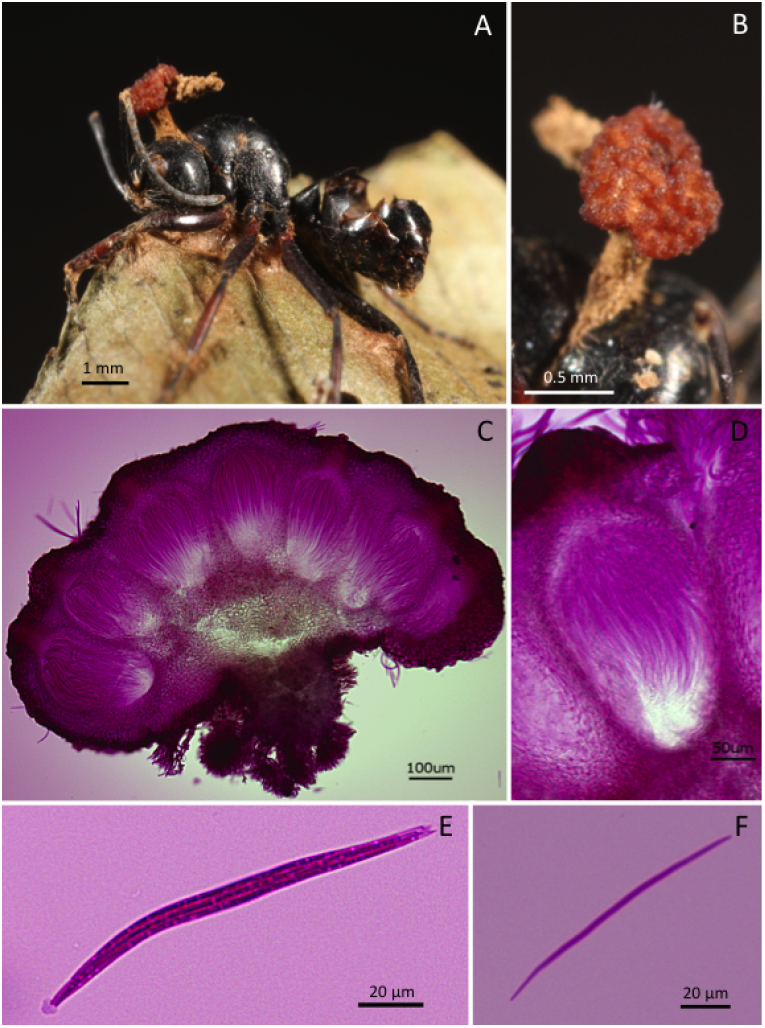
Ophiocordyceps australia (teleomorphic structures). **A)** Polyrhachis sp. biting the edge of a leaf; **B)**Close-up of the orange ascoma

**Fig. 15.**
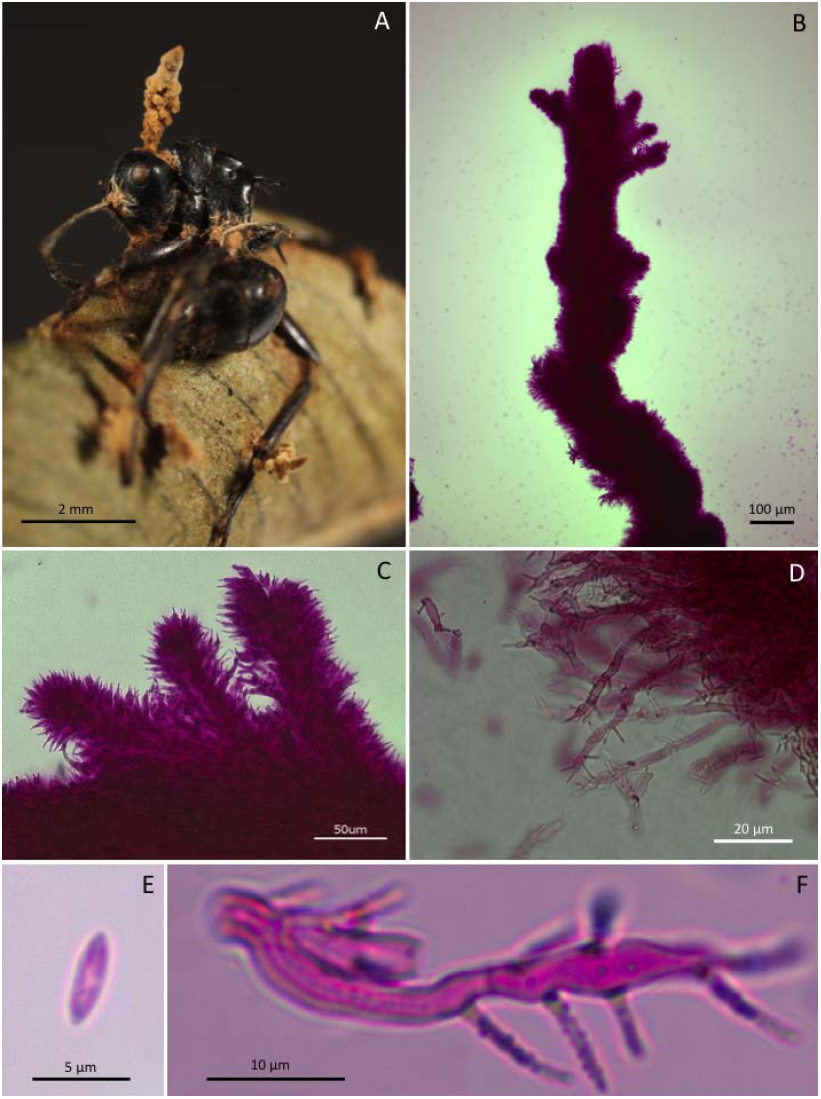
Ophiocordyceps australia (anamorphic structures). **A)** Dead *Polyrhachis* with stromata arising from leg joints and dorsal pronotum. **B)** Synnema **C-D)** Close-up of synnema; **E)** Conidium; **F)** Individual long phialides with multiple verrucose necks.

***Ophiocordyceps polyrhachis-nigrans*** Araújo & D. P. Hughes sp. nov.

External mycelium produced from orifices and sutures; initially white, becoming light-brown with age; Stroma single or branched, produced from dorsal pronotum, averaging 6.5 x 0.3 mm, cylindrical, grayish to light brown; Fertile part produced laterally on the stroma, 1−3, disc-shaped, dark-brown, averaging 1.1 x 0.8 mm. Perithecia immersed to partially erumpent, flask-shaped, 230–260 x 120–150 µm, with short neck. Asci 8-spored, hyaline, cylindrical to clavate, 130–180 x 8–9 µm, cap prominent; Ascospore hyaline, vermiform, straight to gently curved, 85–100 x 3 µm, 5-septate, tapering to both ends.

**Anamorph.** Hirsutella type-A only. Phialides cylindrical to lageniform, 6–8 x 3–4, tapering to a long neck, 9–10 µm long, bearing a terminal conidium, averaging 5x3 µm.

**Germination process.** No germination could be observed since the material examined was not alive.

**Fig. 16.**
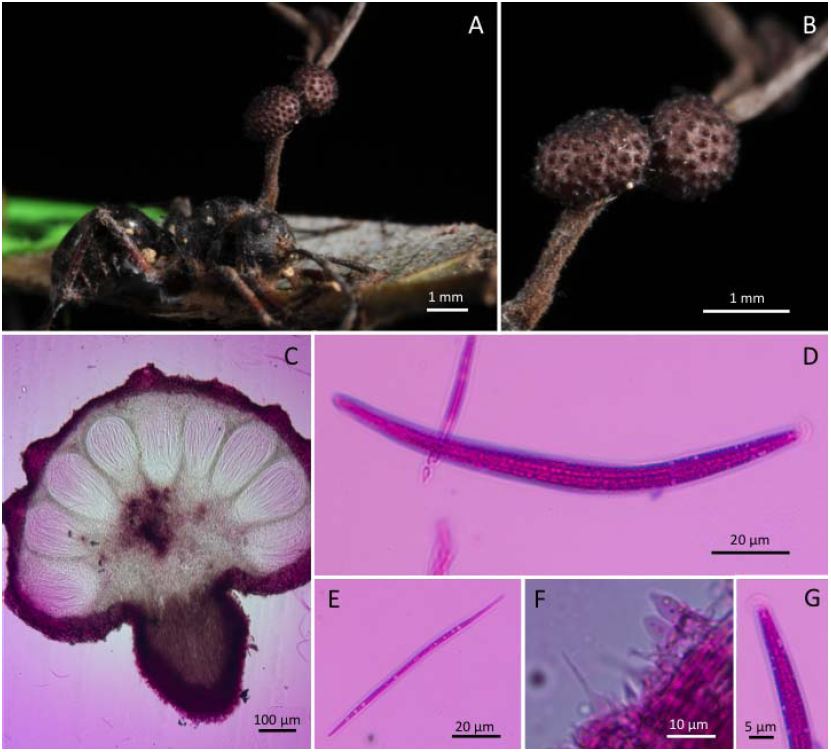
Ophiocordyceps polyrhachis-nigrans. **A)** *Polyrhachis* sp. biting on a leaf edge; B) Close-up showing two ascomatal plates attached to the stroma; **C)** Cross-section of ascoma; **D)** Ascus; **E)** Ascospore; **F)** Hirsutella A-type phialides on stroma; **G)** Ascus-cap close-up.

***Ophiocordyceps clavata.*** (Kobayasi) Araújo & D. P. Hughes, **comb. nov.**

External mycelium scarce, produced mostly on ventral part of the host and mouth. Stromata produced from pronotum, dorsal– and laterally on both sides, clavate, 5–7.5 x 0.35–0.45 (−0.8) mm, never branching. Fertile part produced laterally on one or multiple stromata, 1–6, commonly 2 per stroma, averaging 1 x 0.8 mm, up to 2.5 mmin length; Perithecia immersed to partially erumpent, flask-shaped, 230–270 x 120–160 µm, with short, exposed neck. Asci 8–spored, cylindrical to clavate, 120–160 x 8–10 µm, cap prominent; Ascospore hyaline, cylindrical, straight, rarely curved, 85–100 x 4 µm, 5–septate, one apex roundish, another tapered.

**Anamorph.** Hirsutella type-A only. Cylindrical to lageniform, averaging 12 x 7 µm, tapering to a long neck. No conidia was observed.

**Germination process.** Ascospores germinating in 24h to produce a hair-like 1–3 capilliconidiophores, 40–50 µm long, bearing a terminal, hemispheric capilliconidium, averaging 13 x 3 µm. Some ascospores germinating directly into vegetative hyphae.

**Fig. 17.**
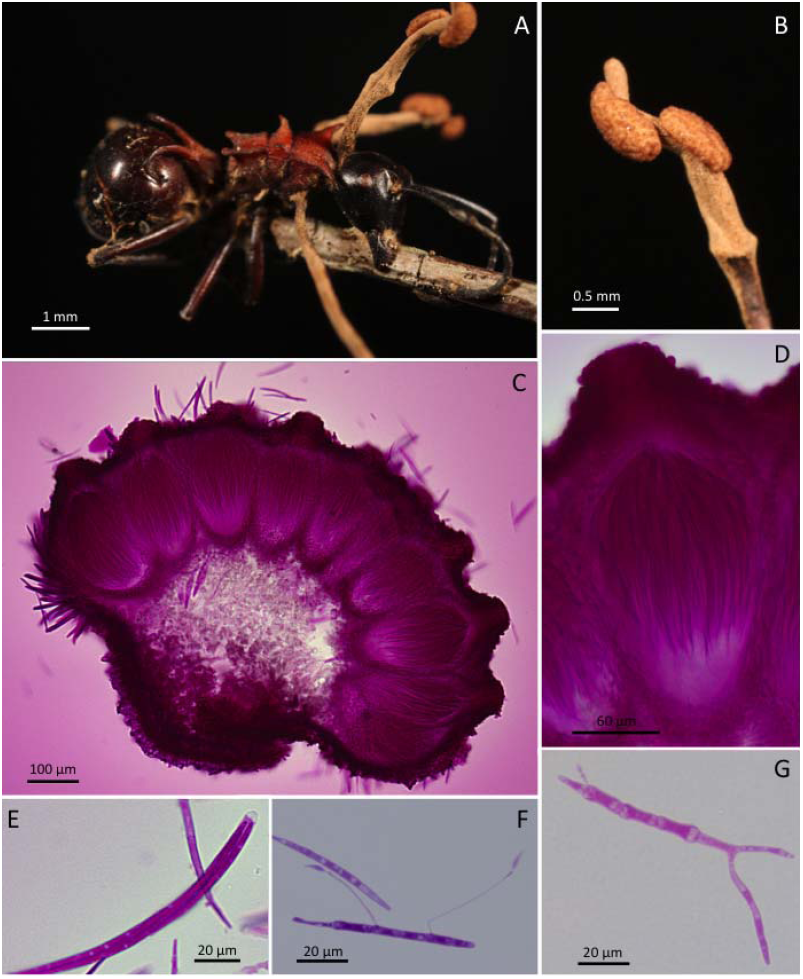
Ophiocordyceps clavata. **A)** *Polyrhachis* sp. with three stromata arising from its body; **B)**Close-up stroma with two ascomatal cushions; **C)**Cross-section of the ascoma, showing the perithecial arrangement; **D)** Close-up perithecium; **E)** Ascus; **F)** Ascospore with two capilliconidiophores bearing one capilliconidium on their apex; **G)** Ascospore germinating on agar plate after 3−5 days.

